# Redox-mediated regulation of a labile, evolutionarily conserved cross-β structure formed by the TDP43 low complexity domain

**DOI:** 10.1101/2019.12.24.885962

**Authors:** Yi Lin, Xiaoming Zhou, Masato Kato, Daifei Liu, Sina Ghaemmaghami, Benjamin P. Tu, Steven L. McKnight

## Abstract

An evolutionarily conserved low complexity (LC) domain is found within a 152 residue segment localized to the carboxyl-terminal region of the TDP43 RNA-binding protein. This TDP43 LC domain contains ten conserved methionine residues. Self-association of this domain leads to the formation of liquid-like droplets composed of labile, cross-β polymers. Exposure of polymers to low concentrations of H_2_O_2_ leads to a phenomenon of droplet melting that can be reversed upon exposure of the oxidized protein to the MsrA and MsrB methionine sulfoxide reductase enzymes, thioredoxin, thioredoxin reductase and NADPH. Morphological features of the cross-β polymers were revealed by a method of H_2_O_2_-mediated footprinting. Similar TDP43 LC domain footprints were observed in highly polymerized, hydrogel samples, liquid-like droplet samples, and living cells. The ability of H_2_O_2_ to impede cross-β polymerization was abrogated by a prominent ALS-causing mutation that changes methionine residue 337 to valine. These observations offer potentially useful insight into the biological role of TDP43 in facilitating synapse-localized translation, as well as aberrant aggregation of the protein in neurodegenerative disease.

## Introduction

A protein designated Tar DNA-binding protein 43 (TDP43) has been the focus of extensive research owing to its propensity to aggregate in the cytoplasm or axoplasm of neurons in patients suffering from neurodegenerative disease (Lee et al., 2011). In a bold and unbiased series of experiments, Lee and colleagues discovered TDP43 aggregates in the brain tissue of disease-bearing patients (Neumann et al., 2006). Over the past decade, TDP43 has emerged as one of the most intensively studied proteins in the field of neurodegenerative disease. The normal biological role of TDP43 has also captured interest owing to its role in the formation of neuronal granules. The TDP43 protein helps facilitate mRNA maturation, mRNA export from nuclei, formation of neuronal granules, and localized, dendritic translation proximal to active synapses (Buratti and Baralle, 2010; Buratti et al., 2001; Chu et al., 2019; Gopal et al., 2017).

TDP43 contains two prototypic RRM domains (Chen-Plotkin et al., 2010), a structured amino-terminal domain that facilitates homotypic oligomerization (Mompean et al., 2016), and a carboxyl-terminal domain of low sequence complexity believed to function either in the absence of structural order (Lim et al., 2016), or via formation of a labile α-helix (Conicella et al., 2016). Human genetic studies of patients suffering from amyotrophic lateral sclerosis (ALS), fronto-temporal dementia (FTD), and other neurodegenerative diseases have led to the discovery of missense mutations clustered within the C-terminal LC domain of TDP43 (Buratti, 2015). It is believed, and in some cases known, that these mutations favor pathological aggregation driven by the TDP43 LC domain. TDP43 aggregation has further been observed in the cytoplasm of neurons under disease conditions driven by expansion of a polyglutamine region of ataxin-2 (Hart and Gitler, 2012), or a hexanucleotide repeat localized within the first intron of the C9orf72 gene (Hsiung et al., 2012).

The LC domains of FUS, many different hnRNP proteins, and many different DEAD box RNA helicase enzymes are typified by the abundant distribution of tyrosine and/or phenylalanine residues. Numerous studies have given evidence that these aromatic residues are important for self-associative interactions that allow for phase transition of LC domains in the form of hydrogels or liquid-like droplets (Kato et al., 2012; Xiang et al., 2015). The LC domain of human TDP43 contains one tyrosine residue and five phenylalanine residues, but is distinguished from prototypic LC domains by the presence of ten evolutionarily conserved methionine residues. Here we show that the TDP43 LC domain self-assembles in a manner specifying a redox-sensitive molecular complex. TDP43 polymers are sensitive to H_2_O_2_-mediated disassembly wherein methionine residues are converted to the methionine sulfoxide state. In this regard, the behavior and biological properties of the TDP43 LC domain are reminiscent of the LC domain specified by the yeast ataxin-2 protein (Kato et al., 2019; Yang et al., 2019). We hypothesize that both the ataxin-2 and TDP43 LC domains assemble into oligomeric structures specifying redox sensors that constitute proximal receptors to the action of reactive oxygen species.

## Results

Recombinant proteins linking the terminal 152 amino acid residues of TDP43 to maltose binding protein (MBP) and/or a 6X-histidine tag were expressed in *E. coli*, purified and incubated under physiologic conditions of monovalent salt and pH (Experimental Procedures). Both fusion proteins were observed to transition into a gel-like state in a manner temporally concordant with the formation of homogeneous polymers as observed by transmission electron microscopy (Figure S1A). Lyophilized gel samples were evaluated by X-ray diffraction, yielding prominent cross-β diffraction rings at 4.7 and 10Å (Figure S1E). When analyzed by semi-denaturing agarose gel electrophoresis, the amyloid-like polymers formed from the TDP43 LC domain dissolved such that the protein migrated in the monomeric state (Figure S1B). As such, we conclude that the TDP43 LC domain is capable of forming labile, cross-β polymers similar to polymers formed by the LC domains of the FUS protein (Kato et al., 2012), various hnRNP proteins (Xiang et al., 2015), and the head domains of various intermediate filament proteins (Lin et al., 2016).

Prior to gelation, solutions containing the 6XHis-tagged LC domain of TDP43 became cloudy. Light microscopic examination of the solutions revealed uniform droplets 2 to 10 μm in diameter. Recognizing that the TDP43 LC domain contains 10 evolutionarily conserved methionine residues, we exposed the observed droplets to varying concentrations of hydrogen peroxide (H_2_O_2_). Droplet melting was initially observed at 0.03% H_2_O_2_, and droplets fully disappeared at 0.3% H_2_O_2_ (Figure 1B). By contrast, no evidence of melting was observed for liquid-like droplets formed from the LC domain of FUS even upon exposure to 1% H_2_O_2_ (Kato et al., 2019). Upon resolving the protein samples variously exposed to H_2_O_2_ by SDS gel electrophoresis, the TDP43 protein was observed to migrate more slowly as a consequence of methionine oxidation (Figure 1A). Direct evidence of H_2_O_2_-mediated methionine oxidation of the TDP43 LC domain was confirmed by mass spectrometry as will be shown subsequently in this report.

**Figure 1.**
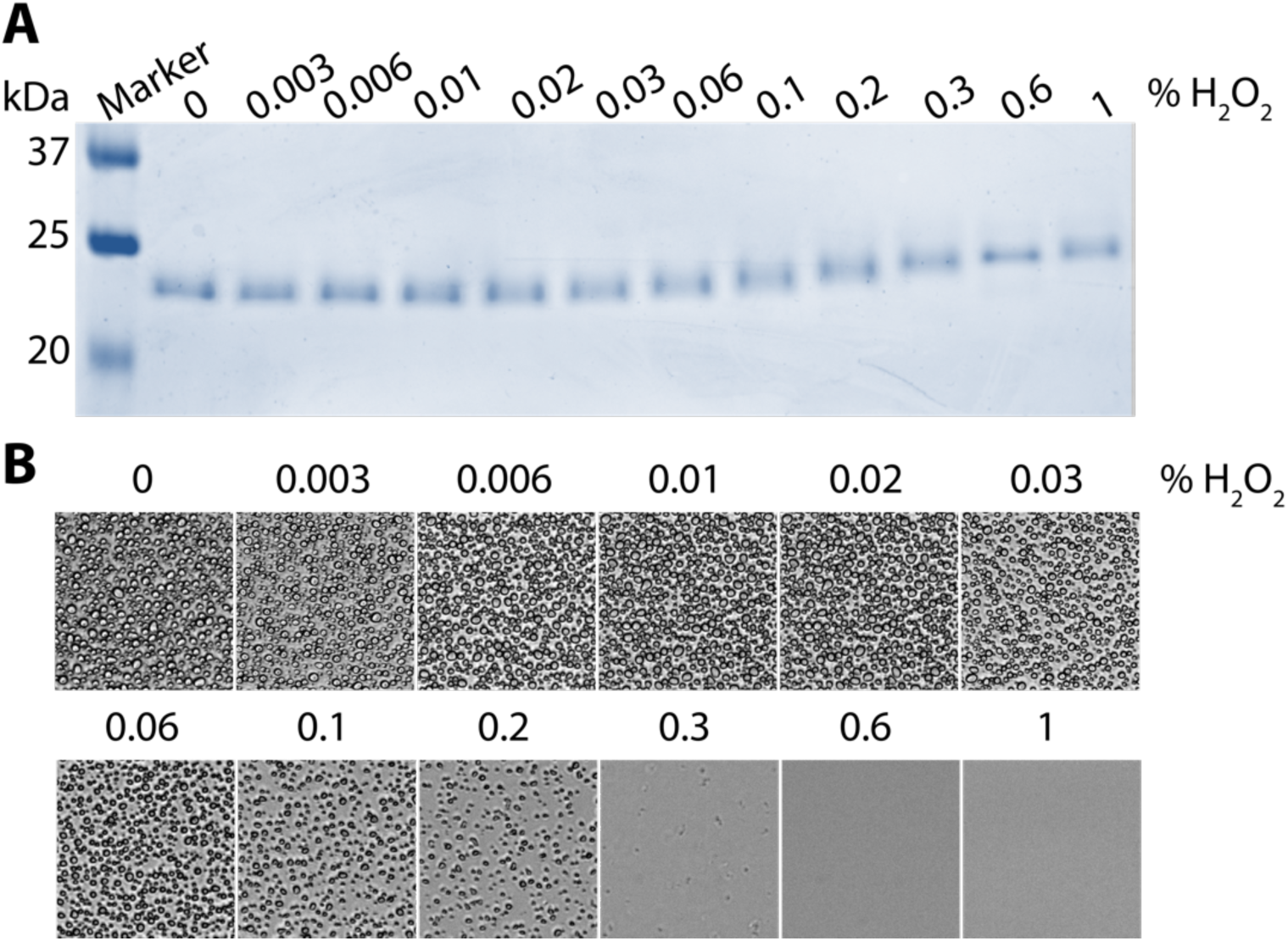
H_2_O_2_-mediated oxidation of the TDP43 low complexity domain. **Top panel (A)** shows Coomassie-stained SDS-PAGE analysis of a purified sample of the TDP43 LC domain after exposure to varied concentrations of H_2_O_2_ ranging from zero % (left) to 1% (right). **Lower panels (B)** show liquid-like droplets formed from the same protein sample used in (A) following exposure for 1 hour to indicated concentrations of H_2_O_2_.

Assuming that liquid-like droplets formed from the TDP43 LC domain might melt as a consequence of methionine oxidation, we tested for droplet re-formation upon addition of the MsrA and MsrB methionine sulfoxide reductase enzymes that are known to reduce the two stereo-isomeric forms of methionine sulfoxide (Moskovitz et al., 1997). H_2_O_2_-mediated oxidation was first quenched by addition of sodium sulfite (Na_2_SO_3_). We then added the MsrA and MsrB enzymes and supplemented the reaction with thioredoxin, thioredoxin reductase and NADPH. These agents allow for sequential steps of reduction of the otherwise oxidized TDP43 methionine residues; reduction of the oxidized MsrA and MsrB enzymes; reduction of oxidized thioredoxin; reduction of oxidized thioredoxin reductase; and terminal conversion of NADPH to NADP^+^. The combination of the five reagents allowed reformation of liquid-like droplets (Figure 2B) and returned the migration pattern of the TDP43 protein to that of the starting protein as deduced by SDS gel electrophoresis (Figure 2A). Removal of either of the two methionine sulfoxide reductase enzymes partially impeded droplet reformation, and partially restored the more rapid migration of the TDP43 substrate protein on SDS gels. By contrast, droplet reformation was fully eliminated upon removable of: (i) both Msr enzymes; (ii) thioredoxin; (iii) thioredoxin reductase; or (iv) NADPH.

**Figure 2.**
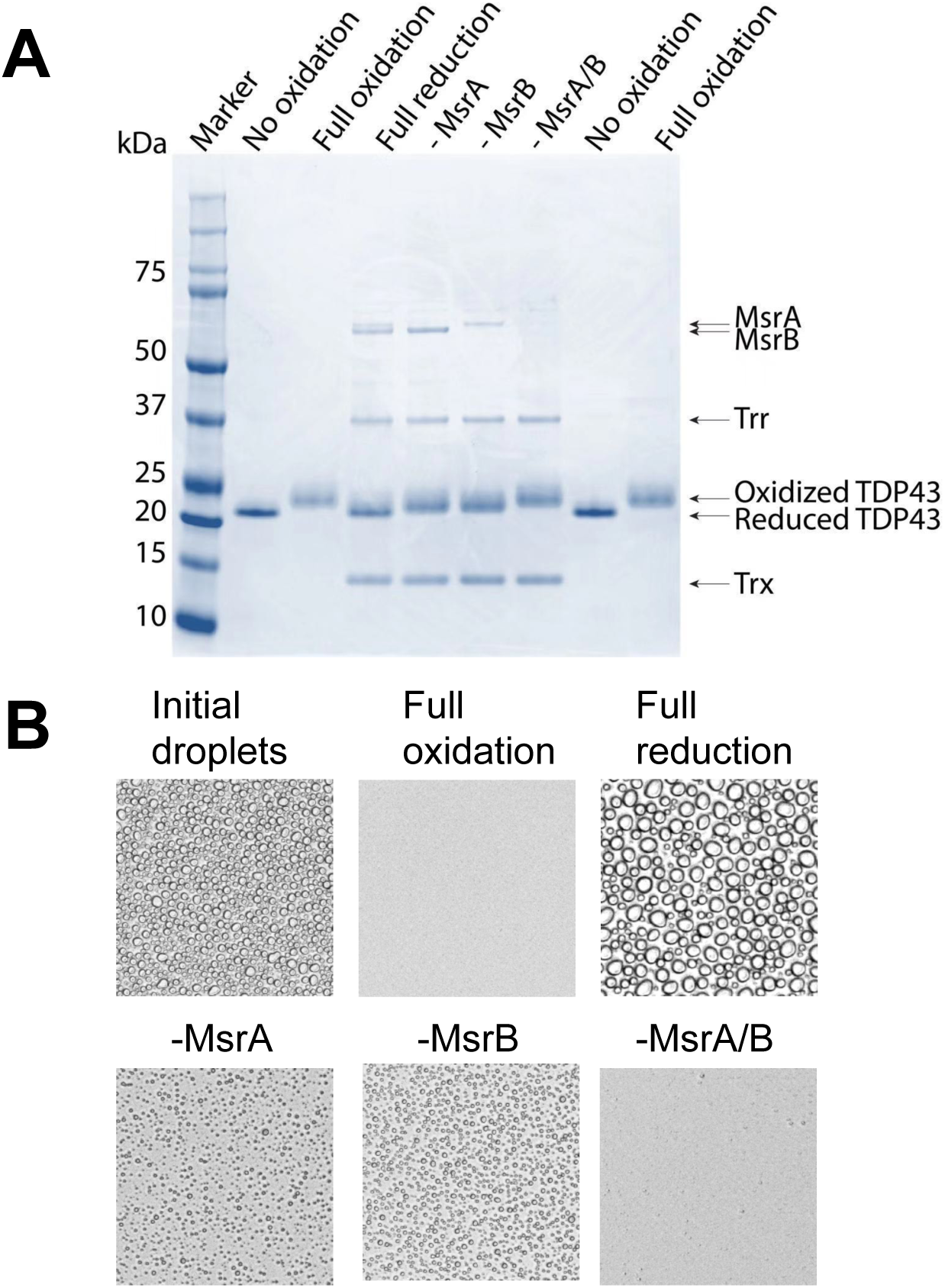
Reduction of the oxidized TDP43 LC domain by two methionine sulfoxide reductase enzymes revives formation of liquid-like droplets. **Top panel (A)** shows an SDS-PAGE gel used to resolve electrophoretic migration patterns of the TDP43 LC domain. H_2_O_2_-mediated oxidation retarded the migration of the TDP43 LC domain (lane 3). Incubation of the oxidized protein with both MsrA and MsrB methionine sulfoxide reductase enzymes, thioredoxin, thioredoxin reductase and NADPH restored the migration pattern to that of the reduced protein (lane 4). Removal of either Msr enzyme led to partial restoration of the electrophoretic mobility of the protein (lanes 5 and 6). Removal of both Msr enzymes failed to alter the electrophoretic pattern of the oxidized protein (lane 7). Lanes 8 and 9 were loaded with fully reduced and fully oxidized proteins, respectively. **Bottom panel (B)** shows photographs of liquid-like droplets formed by un-treated, recombinant TDP43 LC domain (upper left), H_2_O_2_-oxidized protein (upper middle), and H_2_O_2_-oxidized protein exposed to both MsrA and MsrB methionine sulfoxide reductase enzymes, thioredoxin, thioredoxin reductase and NADPH (upper right). Lower three images show droplets formed from enzymatic reduction reactions missing only MsrA (lower left), only MsrB (lower middle), or both methionine sulfoxide reductase enzymes (lower right).

A method of H_2_O_2_-mediated footprinting was used to characterize cross-β polymers formed from the TDP43 LC domain (Figure S2). Following concepts articulated in a recent study of proteome-wide susceptibility to H_2_O_2_-mediated oxidation (Walker et al., 2019), it was reasoned that regions of the TDP43 LC domain directly involved in the formation of structural order might be partially immune to oxidation relative to regions remaining in a state of molecular disorder. Hydrogel preparations of the TDP43 LC domain were exposed for 5 min to a buffer supplemented with varying concentrations of H_2_O_2_. The reaction was quenched by addition of 300mM Na_2_SO_3_, desalted to remove the excess sodium sulfite, denatured in 6M guanidine HCl, and fully oxidized by exposure to 1% ^18^O-labeled H_2_O_2_. The terminal step of saturation oxidation with ^18^O-labeled H_2_O_2_ served two purposes. First, it ensured that no spurious ^16^O oxidation could take place during mass spectrometry sample preparation and analysis. Second, it allowed for accurate quantitation of the fractional oxidation from mass spectrometry data by measuring the ^16^O/^18^O ratio for individual methionine residues (Bettinger et al., 2019). The protein was then fragmented to completion with chymotrypsin and subjected to mass spectrometry as a means of determining the ratio of ^16^O to ^18^O for each of the ten methionine residues resident within the TDP43 LC domain (Experimental Procedures).

We interpret high ^16^O/^18^O ratios equivalent to fully denatured TDP43 to be reflective of unstructured methionine residues, and low ^16^O/^18^O ratios to be reflective of methionine residues that might be involved in formation of cross-β structure. Figures 3 shows a varied pattern of protection for the ten methionine residues localized within the TDP43 LC domain. This pattern corresponds to an H_2_O_2_ “footprint” of the TDP43 LC domain observed in highly polymeric, hydrogel samples of the protein (Figure 3A – left panel). A footprint of protein present in liquid-like droplets is shown in the middle panel of Figure 3A. These similar footprints predict methionine residues M322 and M323 to be substantially protected from H_2_O_2_-mediated oxidation. More modest levels of protection were observed for methionine residues M336, M337 and M339. The two methionine residues located closest to the carboxyl terminus of the protein (M405 and M414), and the two methionine residues located on the amino terminal side of the LC domain (M307 and M311), were poorly protected from H_2_O_2_-mediated oxidation in both hydrogel and liquid-like droplet samples of the TDP43 LC domain.

**Figure 3.**
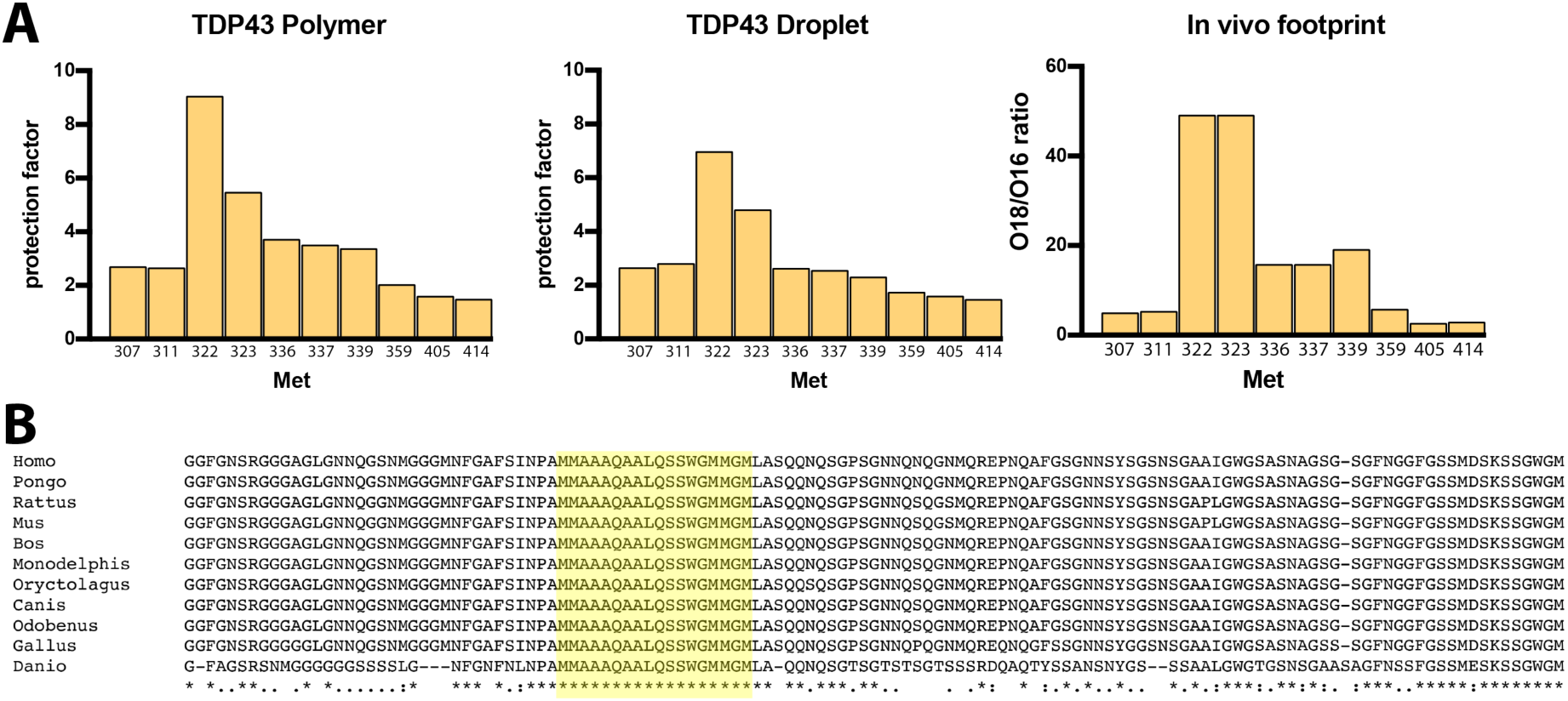
H_2_O_2_-mediated footprints of TDP43 protein present in hydrogel samples, liquid-like droplets and HEK293 cells. For recombinant protein samples used to form hydrogels (left panel) and liquid-like droplets (middle panel), samples were exposed to varying concentrations of H_2_O_2_ for 30 minutes. After quenching reactions with sodium sulfite, samples were denatured in 6M guanidine hydrochloride and exposed for 30 minutes to a 0.5% solution of ^18^O-labeled H_2_O_2_. The ^16^O/^18^O ratio for each methionine residue within the TDP43 LC domain was then determined from chymotrypsin digested samples by mass spectrometry and normalized relative to denatured monomeric protein to determine protection factor calculations (Experimental Procedures). For cellular TDP43 (right panel), HEK293 cells were exposed for 5 minutes to 0.1% H_2_O_2_. Cells were cryo-mill disrupted and solubilized in 4M urea. Flag-tagged TDP43 was recovered by immuno-precipitation, exposed to 5% SDS to fully denature the protein, and exposed for 30 minutes to a 0.5% solution of ^18^O-labeled H_2_O_2_. Following chymotrypsin fragmentation, the 18O/^16^O ratio for each methionine residue was measured by mass spectrometry (Experimental Procedures). For all three samples the region exhibiting most substantial protection from initial ^16^O-H_2_O_2_-mediated oxidation included methionine residues 322, 323, 336, 337 and 339. The location of this footprinted region of 18 amino acids is highlighted in yellow on the sequences of the TDP43 LC domains of eleven vertebrate species ranging from humans (*Homo*) to fish (*Danio*).

Moving from test tube studies of recombinant protein to living cells, we used CRISPR methods to insert both a Flag epitope tag and green fluorescent protein (GFP) onto the amino terminus of the endogenous TDP43 protein of HEK293 cells (Experimental Procedures). Live cell imaging of the GFP-tagged protein revealed punctate nuclear staining (Figure S3) consistent with numerous published studies of TDP43 (Appocher et al., 2017; Wobst et al., 2017). In order to probe for the presence or absence of structural order within the TDP43 LC domain in living cells, we exposed the CRISPR-modified HEK293 cells for 5 minutes to varying levels of H_2_O_2_. The cells were then frozen in liquid nitrogen and powdered using a cryo-mill instrument (Experimental Procedures). Cryo-mill powder was resuspended in lysis buffer and separated into soluble and insoluble fractions by centrifugation. As shown in Figure S4, graded increases in H_2_O_2_ led to graded changes in the ratio of soluble/insoluble TDP43. In the absence of added H_2_O_2_, Western blot analysis revealed that approximately 50% of the cellular TDP43 remained in an insoluble state. This amount of insoluble protein was reduced in a graded manner upon exposure of cells to 0.01%, 0.03% or 0.1% H_2_O_2_, and fully eliminated from cells exposed to either 0.3% or 1% H_2_O_2_. These H_2_O_2_-mediated decreases in the insoluble fraction of TDP43 were reciprocally balanced by increases in the amount of soluble protein.

Recognizing that H_2_O_2_-mediated oxidation of TDP43 might influence its solubility, we resuspended cryo-mill powder in lysis buffer supplemented with 4M urea. Addition of the chaotropic agent allowed for complete solubilization of TDP43 under conditions compatible with immuno-precipitation using anti-Flag antibodies. We thus treated growing cells for 5 minutes with 0.1% of ^16^O-labeled H_2_O_2_, prepared cryo-mill lysate, solubilized the powder with lysis buffer containing 4M urea, and immuno-precipitated the Flag-tagged TDP43 protein. Post-immunoprecipitation the protein was denatured in 5% SDS and exposed to 1% ^18^O-labeled H_2_O_2_ in efforts to saturate oxidation of all methionine residues within the TDP43 LC domain. The sample was quenched with sodium sulfite, desalted, chymotrypsin digested to completion and evaluated by mass spectrometry to measure the ^16^O/^18^O ratio of each of the 10 methionine residues within the TDP43 LC domain. As shown in the right panel of Figure 3A, the oxidation footprint observed for cellular TDP43 was similar to the footprints observed in both hydrogel and liquid-like droplet preparations of the recombinant protein.

The location of the methionine oxidation footprint of TDP43, as defined by the boundaries of methionine residues 322 and 339, is highlighted in yellow upon the sequences of the LC domains of 12 vertebrate species, ranging from fish to humans (Figure 3B). The footprinted region co-localizes with an ultra-conserved segment of 22 amino acids. In the 500 million years separating fish from humans, not a single amino acid has been changed within this ultra-conserved region in any of the 12 species included in this analysis.

The footprinted and ultra-conserved region of the TDP43 LC domain co-localizes with a cross-β structure described from cryo-EM studies recently reported by Eisenberg and colleagues (Cao et al., 2019). Highly related structures were independently resolved for three different cross-β polymers formed from the TDP43 LC domain (Figure 4). Structural overlap among the three independent fibrils was observed to initiate at proline residue 320 of the TDP43 LC domain and persist for 15 residues to tryptophan residue 334. In Figure 4 we compare the methionine oxidation footprint described in this study with the cross-β structures resolved by the Eisenberg lab. The two methionine residues most protected from H_2_O_2_-mediated oxidation, M322 and M323, are disposed wholly within the cross-β structure. The moderately protected methionine residues M336, M337 and M339 are located on the immediate C-terminal side of the cross-β structure. Finally, all five of the oxidation-exposed residues of M307, M311, M359, M405 and M414 were located outside of the Eisenberg cross-β structure.

**Figure 4.**
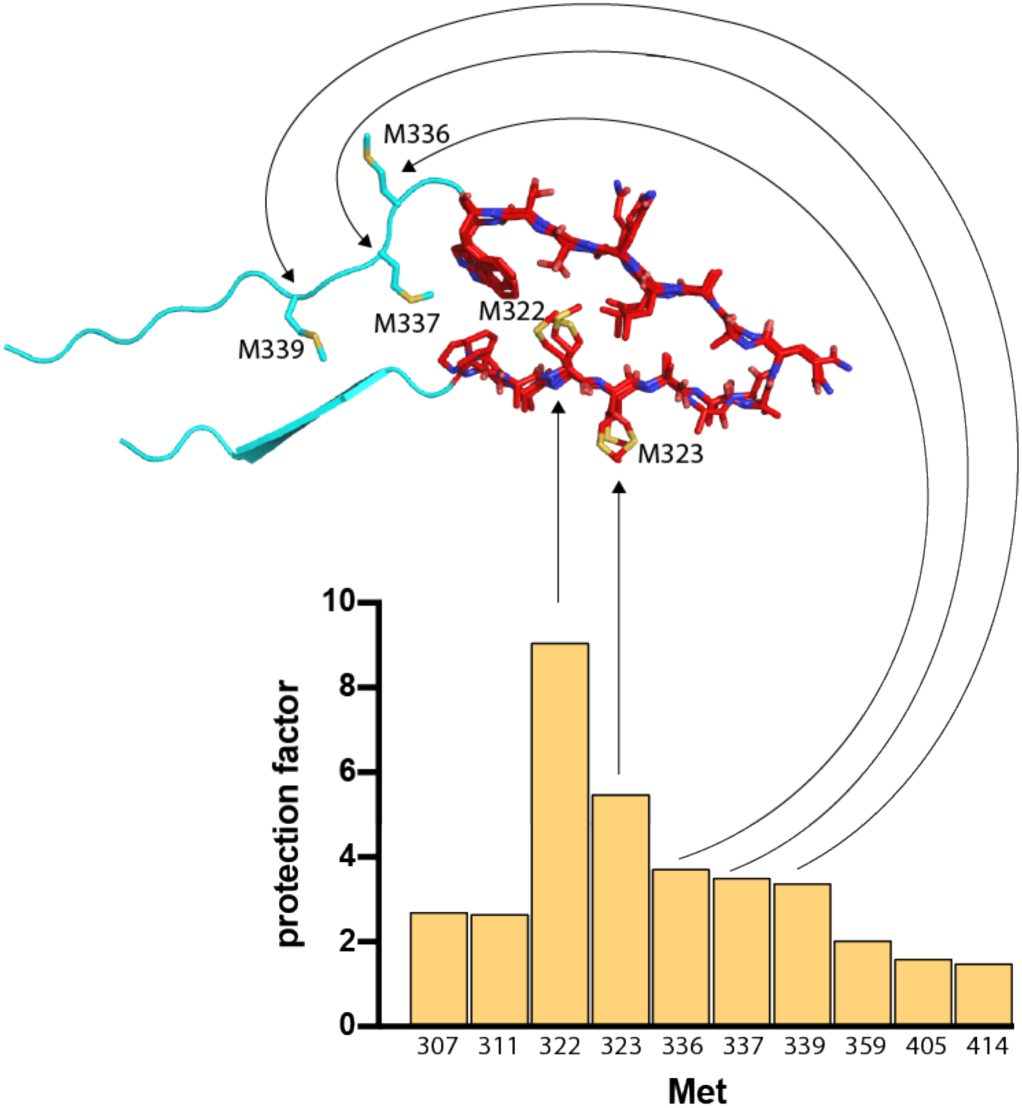
Correspondence of TDP43 LC domain oxidation footprint and molecular structure of cross-β polymers as deduced by cryo-electron microscopy. Arrows connect methionine residues protected from H_2_O_2_-mediated oxidation (TDP43 LC domain hydrogel footprint) with their position either within the cross-β polymer core (M322 and M323), or on the immediate, C-terminal side of the polymer core (M336, M337 and M339). Molecular model of TDP43 LC domain polymer core is derived from cryo-EM structures resolved for three independent TDP43 LC domain fibrils (Cao et al., 2019).

The potential roles of different methionine residues in helping establish the biological utility of the TDP43 LC domain were investigated by introducing methionine-to-valine substitutions into the protein sequence. Each of the ten M-to-V variants was expressed, purified and tested in assays for liquid-like droplet formation and cross-β polymerization as deduced by thioflavin-T staining. Eight of the ten M-to-V variants generated liquid-like droplets indistinguishable from droplets formed from the native LC domain of TDP43 (Figure S5). The M337V variant produced smaller droplets that were noticeably less translucent than those formed by the native LC domain, and the M323V variant yielded amorphous aggregates instead of liquid-like droplets.

The cross-β polymerization capacity of the native LC domain of TDP43 was compared with that of the nine droplet-competent M-to-V variants under three conditions: (i) in the absence of H_2_O_2_-mediated oxidation; (ii) following mild oxidation; and (iii) following extensive oxidation. Partial and full oxidation reactions were carried out in the presence of 6M guanidine such that oxidation would not be influenced by any form of protein structure (Experimental Procedures). After quenching with sodium sulfite, samples were analyzed by SDS-PAGE to ensure equivalent levels of protein integrity and degree of oxidation (Figure 5A). Polymerization was initiated via dilution out of 6M guanidine and monitored by spectrophotometry using thioflavin-T fluorescence (Figure 5B). In the absence of oxidation, all nine variants were observed to polymerize as rapidly as the native LC domain, with mildly enhanced polymerization observed for the M336V, M337V and M339V variants. Under saturating conditions of H_2_O_2_-mediated oxidation, none of the ten proteins displayed any evidence of polymerization. Finally, under conditions of partial oxidation, no polymerization was observed for the native TDP43 LC domain. Mild oxidation similarly prevented polymerization for six of the M-to-V variants. Surprisingly, the M337V variant, and to lesser extents the M336V and M339V variants, polymerized despite partial oxidation.

**Figure 5.**
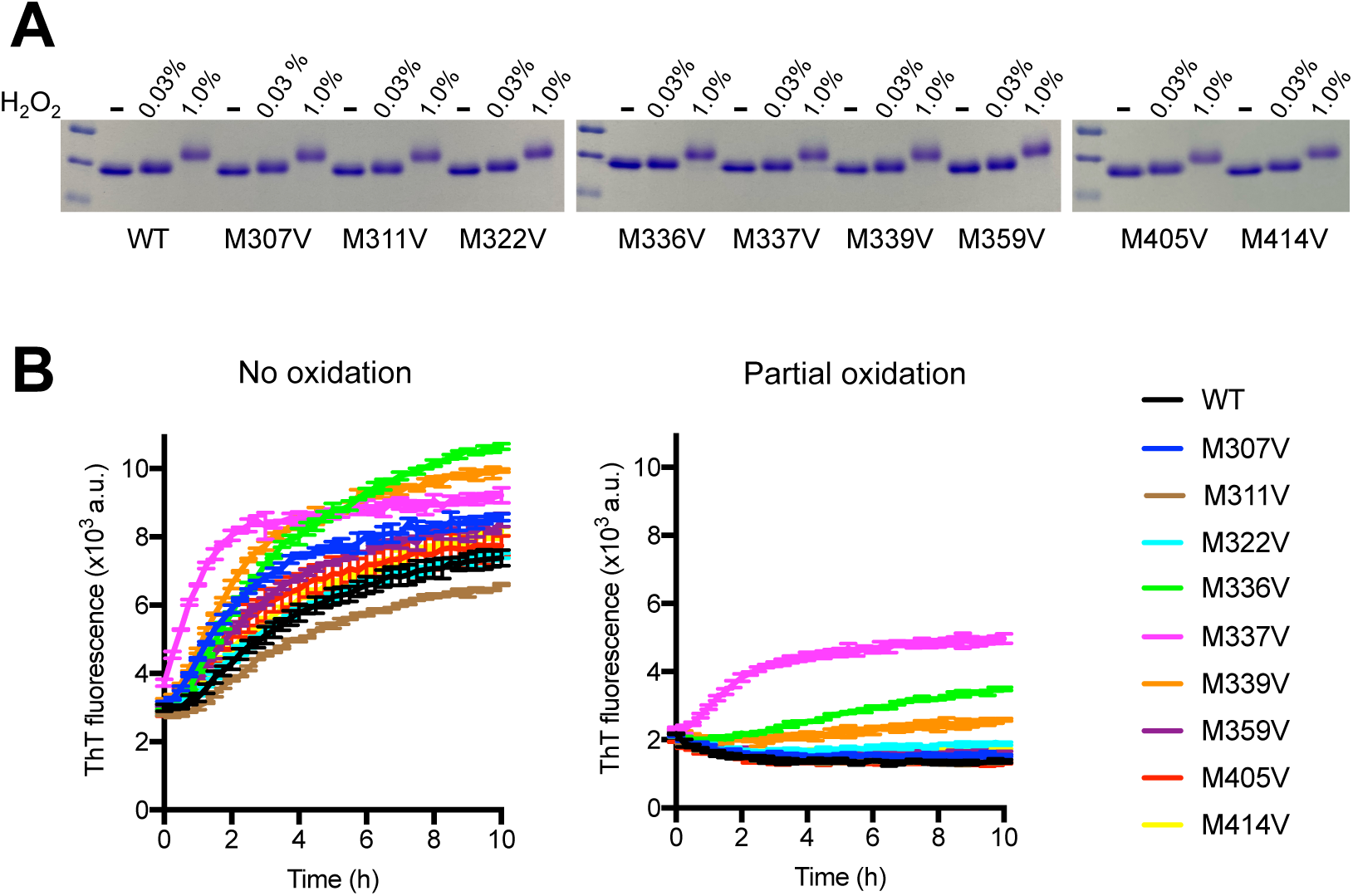
Polymerization assays for nine methionine-to-valine variants within the TDP43 LC domain. Recombinant protein samples were prepared from the native (WT) TDP43 LC domain and nine methionine-to-valine variants. Samples were prepared in the reduced state (-) as well as after exposure for 25 minutes to either 0.03 or 1% H_2_O_2_ in the presence of 6M guanidine HCl. Reduced, partially oxidized, and heavily oxidized samples of each protein were resolved by SDS-PAGE **(Panel A)**. Oxidation reactions were quenched with sodium sulfite, and polymerization was monitored by thioflavin-T (ThT) fluorescence immediately following 20X dilution into polymerization buffer (Experimental Procedures). All ten of the reduced samples exhibited robust evidence of polymerization, with the M337V variant reproducibly exhibiting an enhanced rate of polymerization **(Panel B, left)**. Upon partial oxidation resulting from exposure to 0.03% H_2_O_2_, polymerization was substantially inhibited for all samples save for the M337 variant **(Panel B, right)**. No evidence of polymerization was observed for the native protein or any of the nine methionine-to-valine variants subsequent to exposure to 1% H_2_O_2_ (data not shown).

## Discussion

Here we describe studies of the low complexity domain of the TDP43 protein. We offer the following observations new to the study of this protein. First, the LC domain of TDP43 assembles into a cross-β structure that accounts for localized protection from H_2_O_2_-mediated oxidation. The morphological properties of the observed pattern of oxidation protection yield a footprint that is similar whether taken from highly polymerized, hydrogel samples of the protein, liquid-like droplet samples of the protein, or the TDP43 protein endogenous to living cells.

Second, the location of this footprinted region of the TDP43 LC domain corresponds precisely to a region of extreme evolutionary conservation. The footprinted region is also coincident in location to that of a cross-β polymer core of the TDP43 LC domain characterized at a molecular level by cryo-electron microscopy (Cao et al., 2019). Indeed, the idiosyncratic properties of the TDP43 LC domain footprint correlate with the molecular structure of the cross-β polymers resolved by Eisenberg and colleagues. By contrast, our observations are not easily reconciled with the reported presence of localized α-helical structure within the TDP43 LC domain as deduced by solution NMR spectroscopy (Conicella et al., 2016).

Third, methionine-to-valine mutation of residues 336, 337 and 339 of the TDP43 LC domain yielded proteins that polymerize more rapidly than normal, and are partially resistant to H_2_O_2_-mediated inhibition of polymerization. We make note of the fact a M337V mutation has been observed in independent kindreds by human genetic studies of amyotrophic lateral sclerosis (Rutherford et al., 2008; Tamaoka et al., 2010), and that CRISPR-mediated introduction of this single amino acid change into the endogenous TDP43 gene of mice leads to profound neuropathology (Ebstein et al., 2019; Gordon et al., 2019). We hypothesize that methionine residues 336, 337 and 339 may be of particular importance to a “redox switch” evolutionarily crafted into the LC domain of TDP43. Whereas methionine residues located outside of the cross-β core of the LC domain are far easier to oxidize than those located proximal to the polymer core, we imagine that oxidation of “easy-to-hit” methionine residues may trigger a cascade that loosens the structure – eventually allowing for oxidation of methionine residues 336, 337 and/or 339. Once oxidized within the four amino acid region specifying these “cardinal” methionine residues, we offer that polymerization of the TDP43 LC domain is effectively blocked until the MsrA and MsrB methionine sulfoxide reductase enzymes are able to reduce the protein and revive capacity for LC domain self-association. Extending from these thoughts, it is possible to appreciate the potential hazard of the M337V mutation, as it might aberrantly allow cytoplasmic aggregation of TDP43 under conditions of partial oxidation.

In many regards, the observations articulated herein parallel similar studies of the low complexity domain of the yeast ataxin-2 protein (Kato et al., 2019; Yang et al., 2019). Those studies provided evidence that the 24 methionine residues within the yeast ataxin-2 LC domain engender redox sensitivity via the same mechanisms described herein for TDP43. The biologic utility of the yeast ataxin-2 redox sensor can be understood from comprehensive studies of that system. Yeast ataxin-2 is required for the metabolic state of mitochondria to be properly coupled to the TOR pathway and autophagy (Yang et al., 2019). Since the yeast ataxin-2 protein forms a cloud-like structure surrounding mitochondria, it is sensible to imagine that its methionine-rich LC domain is proximally poised to sense the presence or absence of mitochondria-generated reactive oxygen species. Indeed, detailed experiments have been undertaken to validate the importance of oxidation-mediated changes in the polymeric state of the yeast ataxin-2 LC domain as a key event in coupling the metabolic state of mitochondria to the activity state of TOR and its influence on autophagy (Kato et al., 2019).

Unlike yeast ataxin-2, TDP43 is primarily a nuclear protein. Why, then, would the LC domain of TDP43 be endowed with the apparent capacity to sense reactive oxygen species? We speculate that when TDP43 shuttles out of the nucleus as a part of mRNP complexes housed within RNA granules, oxidation-sensitive self-association of its LC domain might allow for the “unfurling” of RNA granules in the proximity of mitochondria. This might facilitate localized translation of certain mRNAs in the proximity of mitochondria, perhaps useful for optimizing synthesis of mitochondria-destined proteins in the immediate vicinity of mitochondria.

We close with consideration of the prominent role of TDP43 in neuronal granules (Chu et al., 2019; Gopal et al., 2017). Neuronal granules are found in dendritic extensions of neurons where they are understood to facilitate localized translation proximal to active synapses (Kiebler and Bassell, 2006). Knowing that active synapses are mitochondria-rich (Devine and Kittler, 2018), we offer the simplistic idea that the LC domain of TDP43 might suffer enhanced oxidation upon encountering mitochondria-enriched synapses. If so, we further hypothesize that oxidation-induced dissolution of cross-β polymers otherwise adhering TDP43 to neuronal granules and their resident mRNAs might assist in facilitating synapse-proximal translation.

## Supplementary Data Figures

**Figure S1.**
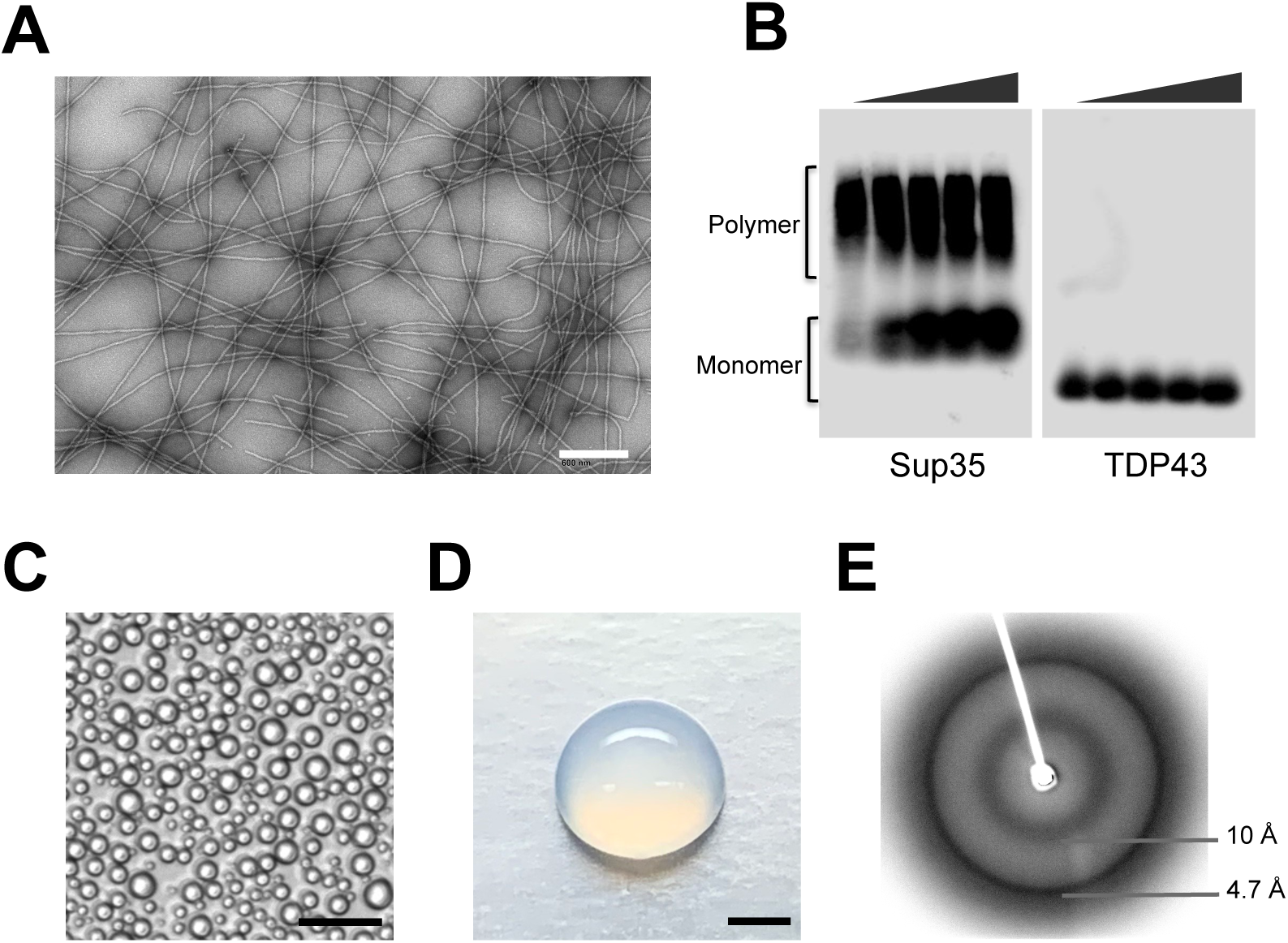
Characterization of labile cross-β polymers formed from the low complexity domain of TDP43. Protein constructs consisting of the LC domain of TDP43 linked to both maltose binding protein (MBP) and a 6Xhistidine tag (6XHis), or the 6XHis tag alone, were expressed in bacterial cells, purified and incubated in gelation buffer to generate uniform polymers (Experimental Procedures). **Panel A** shows a transmission electron micrograph of negatively stained MBP/6XHis-tagged TDP43 LC domain polymers (scale bar = 600 nm). The same polymers were diluted in sample buffer, exposed to varying concentrations of SDS from zero (left lane) to 2% (right lane), electrophoresed on an agarose gel, transferred to nitrocellulose and probed with an antibody to the 6XHis tag (**right side of panel B**). **Left side of panel B** shows irreversibly assembled polymers formed from the yeast Sup35 protein. Liquid-like droplets formed from the 6His-tagged TDP43 protein (**panel C**, scale bar = 25 um) were formed immediately upon dilution of the protein out of the denaturing conditions of 6M guanidine. Clear hydrogel droplets (**panel D**, scale bar = 2 mm) were formed after prolonged incubation of the MBP/6XHis-tagged LC domain of TDP43 in gelation buffer (Experimental Procedures). Hydrogel droplets were buffer-exchanged into distilled water, lyophilized and imaged by X-ray diffraction yielding prominent diffraction rings at 4.7 and 10Å (**panel E**).

**Figure S2.**
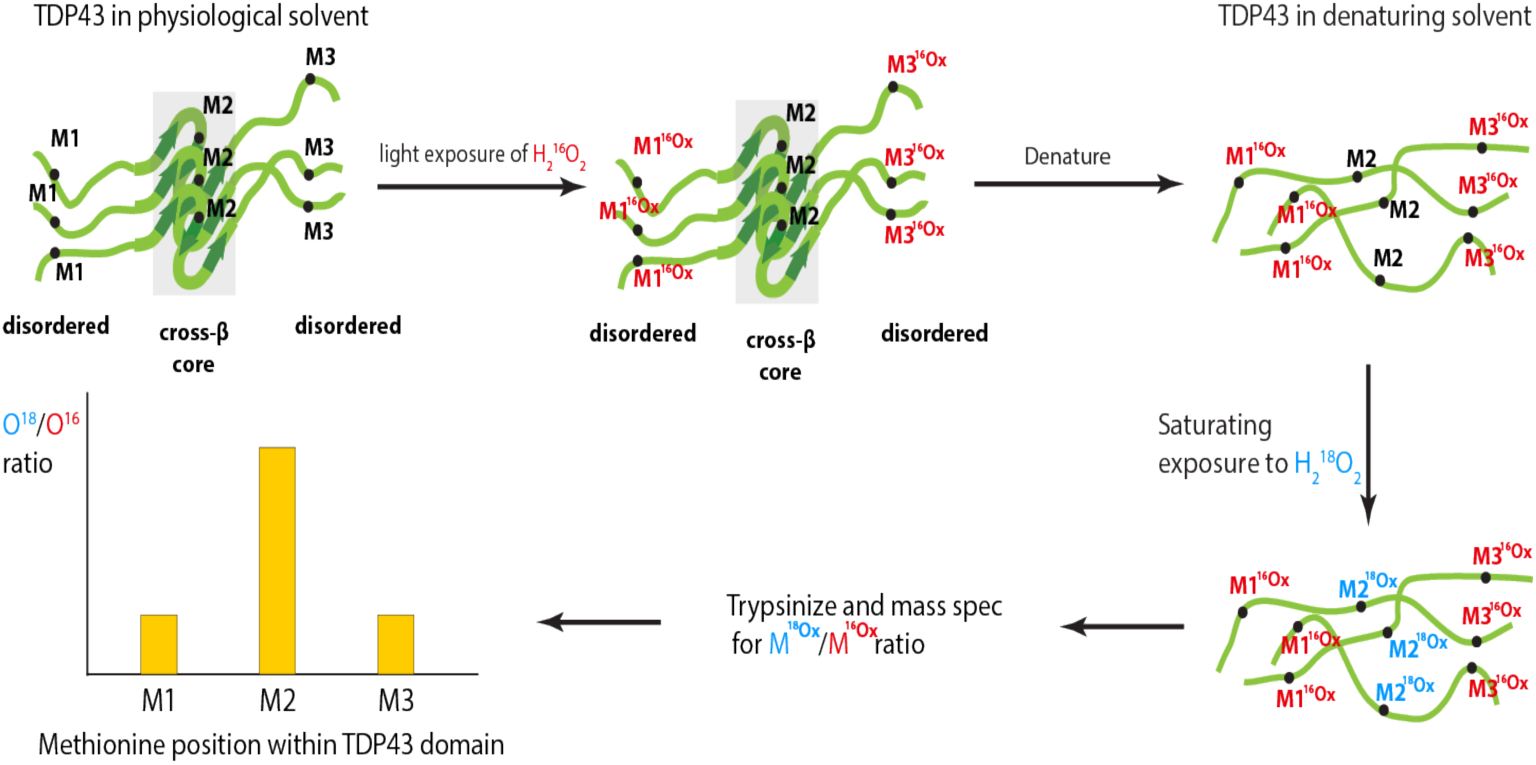
Schematic diagram of methods used for H_2_O_2_-mediated footprinting of TDP43 protein present in hydrogels, liquid-like droplets and living cells. Hypothetical polymers composed of the TDP43 LC domain show a central, cross-β segment (shaded) flanked by regions of structural disorder. Hypothetical methionine residue 2 is localized within cross-β structure; methionine residues 1 and 3 are localized to disordered regions of the protein. Initial treatment with limiting amounts of ^16^O-labled H_2_O_2_ yields oxidation (red color) of methionine residues 1 and 3. Sample is denatured and oxidized to completion with ^18^O-labeled H_2_O_2_, causing oxidation of methionine residue 2 (blue). Samples are then digested with chymotrypsin and analyzed by mass spectrometry to quantitate ^18^O/^16^O ratio of each methionine residue (histogram shown at bottom left).

**Figure S3.**
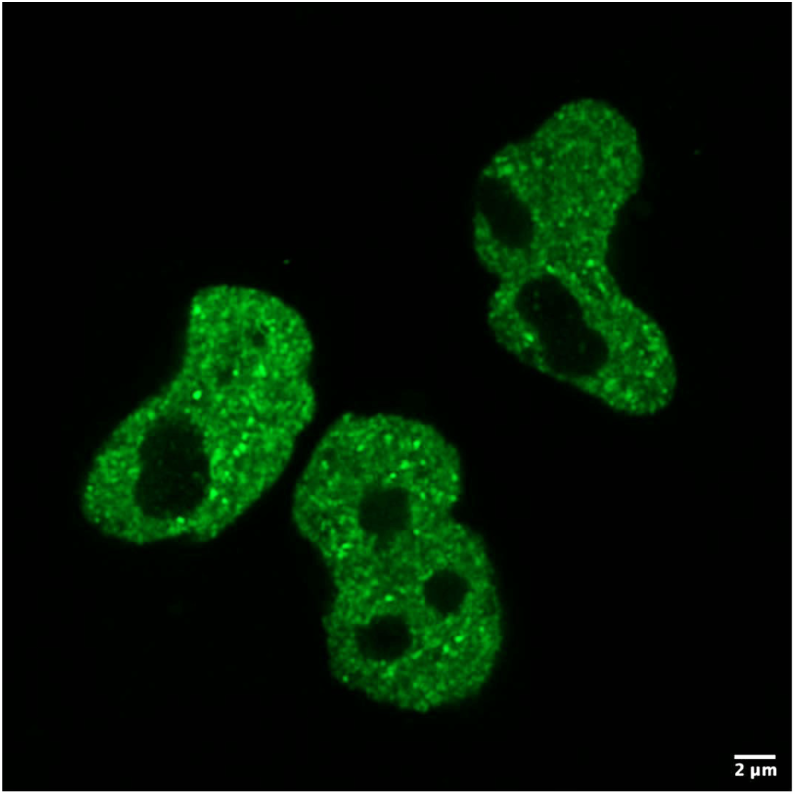
Punctate nuclear distribution of GFP-tagged TDP43 protein in HEK293 cells. CRISPR-tagged TDP43 protein was visualized by live cell imaging (Experimental Procedure). Figure reveals speckled nuclear staining, excluding nucleoli, of endogenous TDP43 protein. Scale bar = 2 microns.

**Figure S4.**
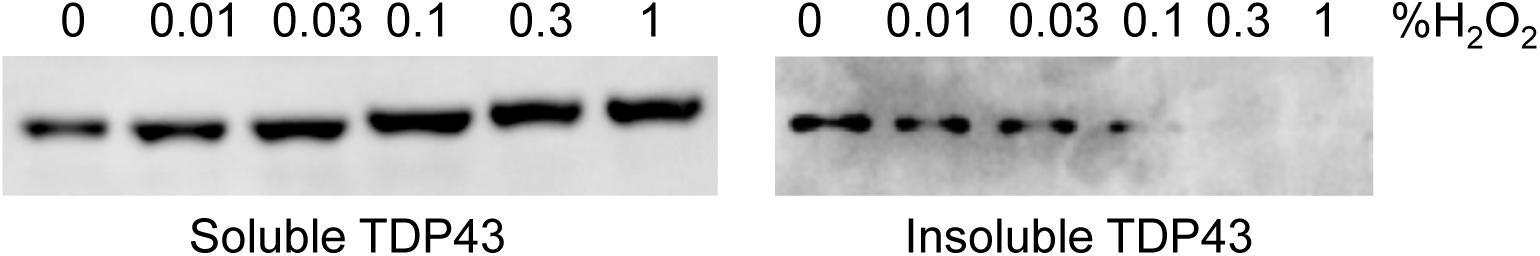
Distribution of cellular TDP43 protein upon lysis of HEK293 cells. The endogenous TDP43 gene of HEK293 cells was CRISPR-tagged at the amino terminus of the protein with both GFP and a Flag epitope (Experimental Procedures). Lysate prepared from cells frozen in liquid nitrogen and disrupted with a cryo-mill instrument was centrifuged to separate soluble from insoluble protein. Protein samples were analyzed by SDS-PAGE and Western blotting using an anti-flag antibody. Unperturbed cells yielded TDP43 protein roughly equally distributed between the soluble **(left panel)** and in soluble **(right panel)** fractions. 5’ treatment of cells with graded increases in the concentration of H_2_O_2_ caused endogenous TDP43 protein to fractionate in a proportionally-increased manner into the soluble fraction of the biochemical lysate. Electrophoretic migration pattern of soluble TDP43 protein **(left panel)** was observed to be retarded in a manner proportional to gradedly increased concentrations of H_2_O_2_.

**Figure S5.**
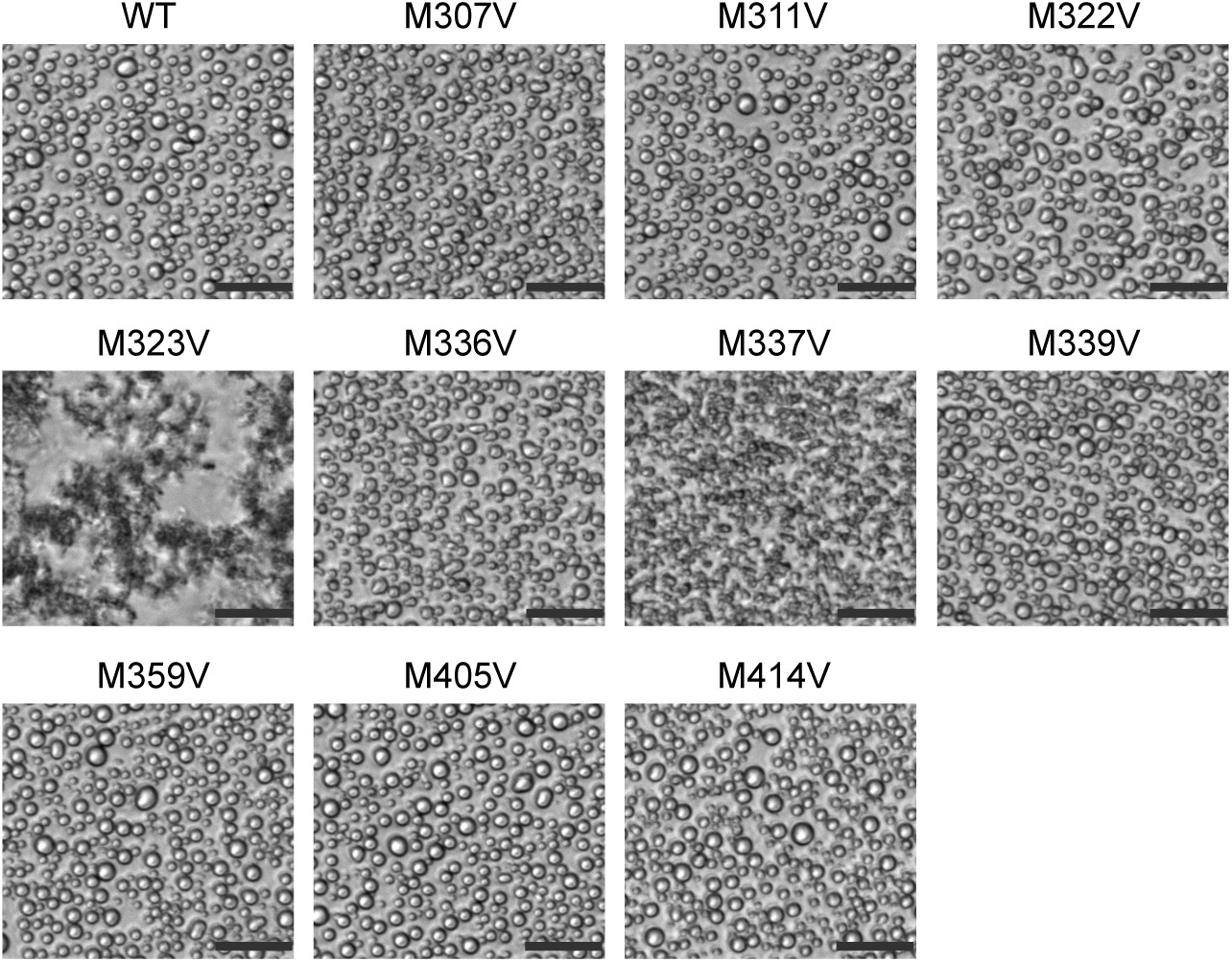
Assays of liquid-like droplet formation of native TDP43 LC domain as compared with variants changing individual methionine residues to valine. Samples of the purified, recombinant 6XHis-tagged TDP43 LC domain were diluted 20-fold into gelation buffer from a stock solution supplemented with 6M guanidine hydrochloride (Experimental Procedures). Liquid-like droplets were formed immediately upon dilution. Indistinguishable droplets were observed for the native protein and eight of the individual methionine-to-valine variants (M307V, M311V, M322V, M336V, M339V, M359V, M405V and M414V). The M323V variant formed amorphous aggregates upon dilution, and the M337V variant formed smaller droplets that were less translucent than those formed by the native protein. Scale bars = 25 μm.

**Figure S6.**
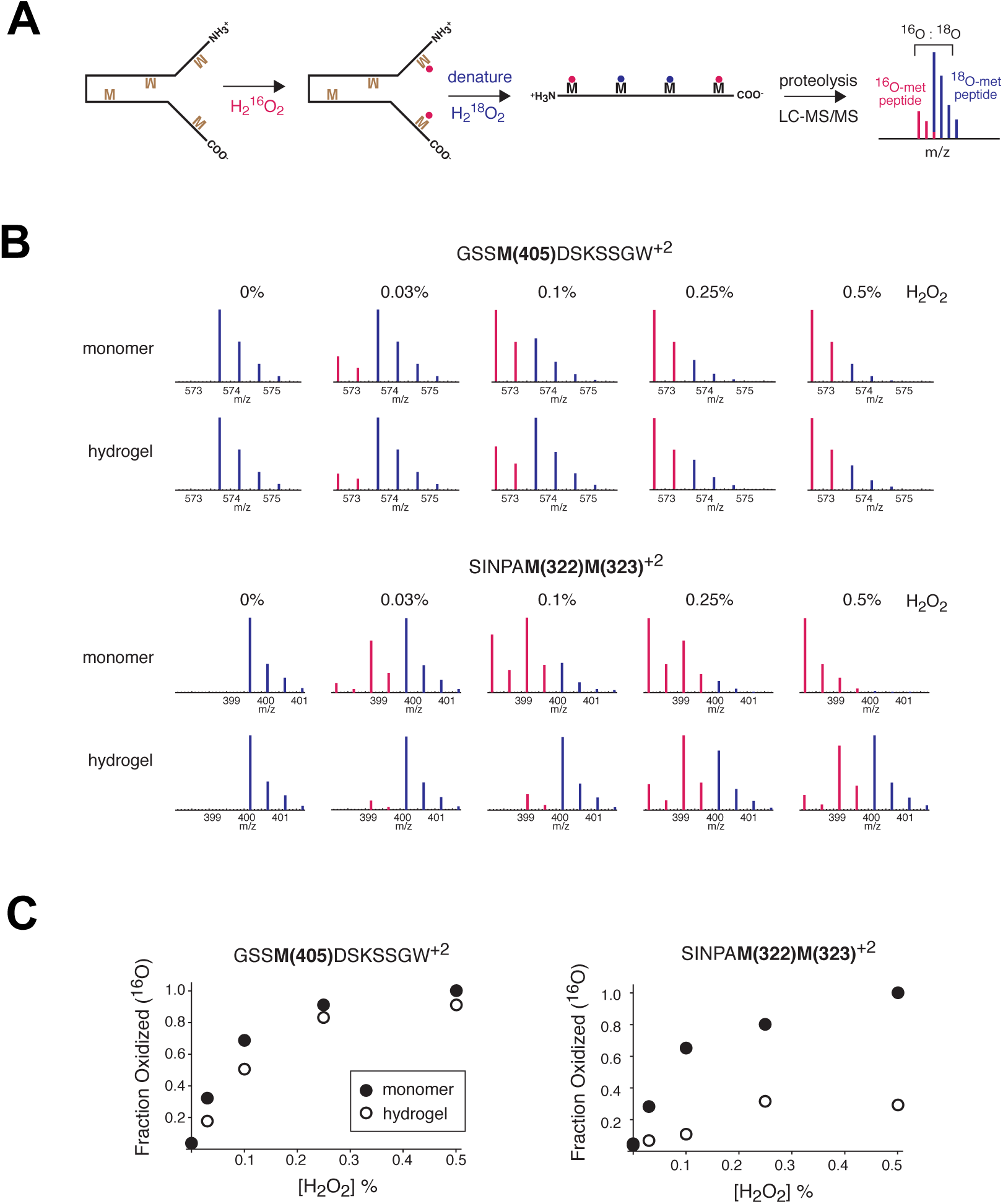
Measurements of fractional oxidation of methionine residues within the TDP43 LC domain as a function of oligomeric or denatured state. **Top panel (A)** shows schematic diagram of partial oxidation of the TDP43 LC domain with ^16^O-H_2_O_2_, followed by denaturation and limit oxidation with ^18^O-H_2_O_2_. **Middle panel (B)** shows MS1 spectra of two peptides located within the TDP43 LC domain. One peptide (above) contained methionine residue 405 that was oxidized equivalently by ^16^O-H_2_O_2_ in the monomeric and hydrogel samples. The other peptide (below) contained methionine residue 323 that was oxidized by ^16^O-H_2_O_2_ more readily in the monomeric state relative to the polymeric, hydrogel state. **Bottom panel (C)** shows measured, fractional levels of ^16^O oxidation for methionine residues 405 (left) and 323 (right) as a function of H_2_O_2_ concentration. Open circles denote protein probed in the polymeric, hydrogel state; closed circles denote protein probed in the denatured state.

## Experimental Procedures

### Cloning of TDP43 constructs

For expression of His-TDP43 LCS in *E. coli*, the TDP43 LC domain (aa 262-414) was amplified using PCR from human cDNA library and cloned into the multiple cloning site of the pHis-parallel1 vector (Sheffield et al., 1999), downstream of T7 promoter. To construct the MBP-TDP43 LCS-His fusion protein, the MBP gene was PCR amplified from pET-His_6_-MBP vector and inserted in pHis-parallel-TDP43 vector and a His_6_ tag was added at the C-terminus. All mutations in TDP43 LCS were introduced via QuikChange site-directed mutagenesis. The organization of all expression vectors and mutated variants was confirmed by DNA sequencing.

### Protein expression and purification

Recombinant proteins were over-expressed in *E. coli* BL21(DE3) cells. The His-TDP43 LC domain was over-expressed with 0.8 mM IPTG in LB medium at 37 °C for 4 hours. The MBP-TDP43 LC domain was over-expressed by induction with 0.6 mM IPTG at 16 °C overnight. For purification of His-TDP43, cells were harvested and resuspended in buffer containing 25 mM Tris-HCl pH 7.5, 200 mM NaCl, 10 mM β-mercaptoethanol (βME) and protease inhibitor and lysed by sonication for 3 minutes (10s on/30s off). His-TDP43 LCD inclusion bodies were collected by centrifugation at 3000 x g for 20 min. The inclusion bodies were dissolved in lysis buffer containing 25 mM Tris-HCl pH7.5, 200 mM NaCl, 6 M guanidine hydrochloride, 10 mM βME and a protease inhibitor tablet (Sigma), followed by sonication for 1 minute (10s on/30s off). The cell lysate was centrifuged at 36,000 x g for 50 minutes. The supernatant was purified through Ni^2+^-NTA resin (Qiagen), washed with buffer containing 25 mM Tris-HCl pH7.5, 200 mM NaCl, 6 M guanidine hydrochloride, 10 mM βME and 20 mM imidazole. Bound protein was eluted from the resin with an elution buffer containing 25 mM Tris-HCl pH7.5, 200 mM NaCl, 6 M guanidine hydrochloride, 10 mM βME and 300 mM imidazole. Purified proteins were concentrated using Amicon Ultra filters (Millipore). Protein solutions were stored at -80°C. Protein purity was confirmed by SDS-PAGE and the concentrations were determined by absorbance at UV_280_.

For purification of the MBP-TDP43-His fusion protein, cells were resuspended and lysed in buffer containing 25 mM Tris-HCl pH7.5, 200 mM NaCl, 2 M urea, 10 mM βME and a protease inhibitor tablet (Sigma), loaded onto Ni^2+^-NTA beads, washed and eluted with similar buffer as His-TDP43 except for addition of 2M urea. The proteins eluted from Ni-NTA column were purified subsequently by amylose resin, eluted in buffer containing 25 mM Tris-HCl pH7.5, 200 mM NaCl, 2 M Urea, 10 mM βME and 10 mM maltose. Proteins were then concentrated using Amicon Ultra centrifugal filters (Millipore), with protein purity confirmed by SDS-PAGE and concentration determination by spectrophometry.

### Phase-separated droplet formation

Phase-separated droplets of His-TDP43 LCD were formed by a quick dilution of the purified protein out of denaturing conditions into gelation buffer containing 25 mM Tris pH 7.5, 200 mM NaCl, 10 mM βME and 0.5 mM EDTA, to reach the final protein concentration of 10-20 μM. Images of liquid-like droplets were taken using Bio-Rad ZOE Fluorescent Cell Imager.

### H_2_O_2_-mediated melting of phase-separated liquid-like droplets

Liquid-like droplet solutions were incubated for 30 min at room temperature. H_2_O_2_ was added to the droplet solution to obtain final concentrations of 0.003%, 0.006%, 0.01%, 0.02%, 0.03%, 0.06%, 0.1%, 0.2%, 0.3% 0.6%, and 1%. After 1 hr incubation at room temperature, droplets images were taken using a Bio-Rad ZOE Fluorescent Cell Imager. To visualize protein oxidation caused by H_2_O_2_, a portion of the reaction mixture was recovered and analyzed by SDS-PAGE.

### Enzymatic reduction of H_2_O_2_-oxidized protein by methionine sulfoxide reductase enzymes

Liquid-like droplets of His-tagged TDP43 LCD (100 μL) were first formed as described above. Solutions were transferred into a 96-well plate and droplets melted by adding H_2_O_2_ to a final concentration of 0.5%. After droplets completely disappeared, 3 μL of 2 M Na_2_SO_3_ was added to the mixture in order to quench residual H_2_O_2_. After incubation for 1 hr, 2.5 mM NADPH solution was added. Subsequently, 10 μL of a 10x enzyme cocktail mix that contained 1 μM MBP-MsrA, 10 μM MBP-MsrB, 100 μM His-tagged Trx1 and 20 μM His-tagged Trr1 was added. The reaction mixture was incubated at room temperature and inspected for droplet revival. Images were taken with Bio-Rad ZOE Fluorescent Cell Imager after incubation overnight. To visualize protein reduction, a portion of the reaction mixture was recovered from the well and analyzed by SDS-PAGE.

### X-Ray diffraction

For X-ray diffraction, His-TDP43 LCD was diluted in gelation buffer at 2 mg/ml. The mixture was incubated for 3 days at 4°C. Polymer pellets were collected by centrifugation at 20,000 x g for 30 min. Pellets were resuspended in 0.2 mL milli-Q water and dialyzed in 1 L milli-Q water for 12 hr twice. Dialyzed samples were lyophilized overnight and exposed to an X-ray beam to obtain diffraction patterns as described previously (Kato et al., 2012).

### Semi-denaturing detergent agarose gel electrophoresis (SDD-AGE)

The stability of cross-β polymers formed from the MBP-TDP43 low complexity domain was tested by semi-denaturing detergent agarose gel electrophoresis (SDD-AGE) as described previously (Kato et al., 2012). Briefly, polymers made from MBP-TDP43 LCD-6XHis and the yeast Sup35NM protein were diluted in gelation buffer at 0.2 mg/ml and 0.1 mg/ml, respectively, sonicated briefly and incubated for 10 min in gelation buffer containing indicated concentrations of SDS (0 - 2%) at 37 °C. Reaction mixtures were then loaded onto a 1.5% agarose gel to separate polymers and monomers. Proteins were transferred onto a nitrocellulose membrane and analyzed by western blotting using a His-tag antibody (Kato et al., 2012).

### Mammalian cell culture and generation of HEK 293 cells harboring CRISPR-edited TDP43

Mammalian cell culture experiments were performed using the HEK293 cell line (ATCC), grown in Dulbecco’s Modified Eagle Medium (Thermo Fisher Scientific), supplemented with 10% fetal bovine serum at 37 °C with 5% CO_2_. For site-specific integration of the 3xFlag-GFP into the endogenous TDP43 locus, three sgRNAs were designed using published methods (Ran et al., 2013). All sgRNAs were designed to target regions close to the TDP43 start codon, and cloned into pGuide-it vector (TaKaRa). To construct the donor vector, ∼1000 bp homologous arms located upstream or downstream from the TDP43 start codon were amplified by PCR. The 3xFlag-GFP sequence was also PCR amplified. The three DNA pieces, upstream of TDP43 start codon, 3xFlag-GFP and downstream were introduced into pGEM-T Easy vector by In-Fusion cloning (TaKaRa). HEK293 cells were co-transfected with the donor vector and sgRNAs using Lipofectamine 2000 (Thermo Fisher Scientific). Post-transfection, cells were split at low density to allow formation of single colonies. GFP-positive colonies containing properly inserted 3xFlag-GFP were screened by anti-Flag western blotting and confirmed by DNA sequencing.

### In vitro footprinting

Three protein samples were prepared: (i) the TDP43 LC domain in a highly polymeric, hydrogel form; (ii) the TDP43 LC domain in the liquid-like droplet form; and (iii) the TDP43 LC domain in the monomeric form (denatured in 6 M guanidine HCl). To obtain TDP43 LCD polymers, His-TDP43 LCD purified in 6 M guanidine buffer was diluted to 100 μM in gelation buffer containing 25 mM Tris pH 7.5, 200 mM NaCl, 10 mM βME and 0.5 mM EDTA, and left at room temperature for two days. For the denatured sample, purified His-tagged TDP43 LCD was dialyzed in denaturing buffer containing 25 mM Tris pH7.5, 200 mM NaCl, 6 M guanidine hydrochloride, 10 mM fresh βME, and concentrated to 100 μM. For liquid-like droplet formation, purified His-TDP43 LCD in 6 M guanidine buffer was diluted to 100 μM in gelation buffer leading to immediate formation of liquid-like droplets.

All samples were oxidized by ^16^O-labeled H_2_O_2_ at concentrations of 0%, 0.03%, 0.1%, 0.25% and 0.5% for 30 min. Oxidation reactions were quenched by addition of twice the molar ratio of Na_2_SO_3_ relative to H_2_O_2_. All samples were then denatured in buffer containing 25 mM Tris pH7.5, 200 mM NaCl, 6 M guanidine hydrochloride, 10 mM fresh βME. To remove H_2_O_2_ and Na_2_SO_3_, buffer was exchanged using a centrifugal filter (Millipore). To achieve full oxidation, 0.5 % ^18^O-labeled H_2_O_2_ was added to the samples, allowing further oxidation for 30 min. Reactions were quenched by addition of twice the molar ratio of Na_2_SO_3_, then processed for mass spectrometry.

### In vivo footprinting

Ten 15 cm dishes of HEK293 3xFlag-GFP CRISPR cells were prepared for each assay point. At ∼80% confluency, cells were treated with four concentrations of ^16^O-labeled H_2_O_2_, 0%, 0.01%, 0.03%, 0.1% for 5 minutes at 37 °C. Cells were harvested, frozen in liquid nitrogen, ground by cryomill (Restch), and resuspended in IP buffer (20 mM Hepes pH7.4, 100 mM Na_2_PO_4_, 50 mM NaCitrate, 100 mM NaCl, 0.1% Tween20, 4 M urea). After brief sonication, the cell lysate was centrifuged at 13,400 rpm for 10 mins at 4 °C. Supernatant was diluted 5-fold in lysis buffer without urea (20 mM Hepes pH7.4, 100 mM Na_2_PO_4_, 50 mM NaCitrate, 100 mM NaCl, 0.1% Tween20), followed by anti-Flag-immunoprecipitation. Cell lysate was bound to Flag-beads, washed three times with IP buffer, and eluted at 95 °C in buffer supplemented with 5% SDS. A solution composed of elution buffer supplemented with 0.5% ^18^O-H_2_O_2_ was added to each sample to effect full oxidation and after 30 minutes quenched with twice the molar ratio of Na_2_SO_3_. Buffer exchange was then performed to move samples into a solution compatible with mass spectrometry (50 mM TEAB, 5% SDS).

### Thioflavin-T assays

The native His-TDP43 LC domain and its methionine-to-valine variants were diluted to 300 μM with buffer containing 25 mM Tris-HCl, pH 7.5, 200 mM NaCl, 6 M guanidine hydrochloride, and treated with buffer (no oxidation), 0.03125% (partial oxidation) and 1.0% (heavy oxidation) H_2_O_2_ for 25 minutes respectively. 200 mM Na_2_SO_3_ was added to quench unreacted H_2_O_2_. The resulting H_2_O_2_-treated samples were diluted to 15 μM with gelation buffer (25 mM Tris-HCl pH 7.5, 200 mM NaCl, 1 mM βME, 30 mM Thioflavin T) and mixed by vortexing. Diluted samples were transferred to a 96 well fluorescence assay plate (Costar assay plate 96, CORNING), at 150 μL per well. The plate was sealed with a foil film to prevent evaporation. Thioflavin-T signal was monitored continuously at 15 min intervals with 20 seconds shaking being applied before each reading. Gelation buffer alone was used as a reading blank.

### Mass spectrometry sample preparation of recombinant proteins

After exposure to ^16^O-labeled H_2_O_2_, complete oxidation by ^18^O-labeled H_2_O_2_, and quenching, each sample consisted of ∼120 μM protein in a buffer consisting of 25 mM Tris pH 7.5, 200 mM NaCl, 6 M guanidinium hydrochloride, 10 mM βME and 163 mM Na_2_SO_3_. Each sample was diluted 10-fold in 25 μL of 10 mM ammonium bicarbonate. Samples were reduced by addition of 25 mM dithiothreitol (DTT) to a final concentration of 2 mM followed by incubation at 55 ^o^C for 1 hour. Iodoacetamide was added to a final concentration of 10 mM and samples were incubated in the dark at room temperature for 30 minutes to alkylate proteins. 1.1 μL DTT was added to quench each reaction. 2 μL of chymotrypsin was added to each reaction to a final concentration of 500 ng/μL and samples were incubated overnight at 37 ^o^C. Formic acid was added to a final concentration of 1%, samples were dried down in a Speed Vac, and subsequently re-suspended in 0.1% trifluoroacetic acid prior to LC-MS/MS analysis.

### Mass spectrometry sample preparation of immuno-purified proteins

Samples were diluted 1:1 in 5% SDS and 50 mM triethylammonium bicarbonate (TEAB) to a final volume of 100 μL. Samples were then reduced with 2 mM DTT, followed by incubation at 55°C for 60 minutes. Iodoacetamide was added to a final concentration of 10 mM and samples were incubated in the dark at room temperature for 30 minutes to alkylate proteins. Phosphoric acid was added to 1.2%, followed by six volumes of 90% methanol and 100 mM TEAB. The entirety of the resulting solution was added to S-Trap micro-spin columns (Protifi) and the columns were centrifuged at 4,000 x g for 1 minute. The S-Traps containing trapped proteins were washed twice by centrifuging through 90% methanol and 100 mM TEAB. 20 μL of 50 ng/μL chymotrypsin was added to each S-Trap and samples were placed in a humidity chamber at 37°C overnight. Subsequently, the S-Traps were centrifuged at 4,000 x g for 1 minute to collect digested peptides. Sequential additions of 0.1% TFA in acetonitrile and 0.1% TFA in 50% acetonitrile were added to the S-trap, centrifuged, and pooled. Samples were frozen and dried down in a Speed Vac, then re-suspended in 0.1% trifluoroacetic acid prior to LC-MS/MS analysis.

### LC-MS/MS methodology

Peptides were injected onto a homemade 30 cm C18 column with 1.8 um beads (Sepax), with an Easy nLC-1200 HPLC (Thermo Fisher), connected to a Fusion Lumos Tribrid mass spectrometer (Thermo Fisher). Solvent A was 0.1% formic acid in water, while solvent B was 0.1% formic acid in 80% acetonitrile. Ions were introduced to the mass spectrometer using a Nanospray Flex source operating at 2 kV. The gradient began at 3% B and held for 2 minutes, increased to 38% B over 15 minutes, then ramped up to 90% B in 2 minutes and was held for 2 minutes, before returning to starting conditions in 2 minutes and re-equilibrating for 7 minutes, for a total run time of 30 minutes. The Fusion Lumos was operated in data-dependent mode, with MS1 scans acquired in the Orbitrap, and MS2 scans acquired in the ion trap. A top-N experiment was used, with the N set to 8. Monoisotopic Precursor Selection (MIPS) was set to Peptide. The full scan was done over a range of 375-1400 m/z, with a resolution of 120,000 at m/z of 200, an AGC target of 4e5, and a maximum injection time of 50 ms. Peptides with a charge state between 2 and 5 were picked for fragmentation. Precursor ions were fragmented by higher-energy collisional dissociation (HCD) using a collision energy of 30%, using an isolation width of 1.1 Da. The AGC was set to 5e4 with a maximum injection time of 240 ms. Dynamic exclusion was set to 20 seconds.

### Measurement of fractional oxidation

Raw files for all samples were searched against the sequence of specific constructs using the integrated Andromeda search engine with MaxQuant software (Cox and Mann, 2008). Peptide and protein quantification were performed with MaxQuant using the default parameter settings and chymotrypsin was selected as the enzyme allowing up to 5 missed cleavages. ^18^O methionine sulfoxide, ^16^O methionine sulfoxide and N-terminal acetylation were set as variable modifications and carbamidomethyl cysteine was set as a fixed modification. Raw files were converted to mzXML format with ProteoWizard’s MSConvert software using the vendor supplied peak picking algorithm and the threshold peak filter set to the top 2000 peaks in each scan (Kessner et al., 2008). The MaxQuant supplied evidence file and the mzXML file were used as input into a custom algorithm, essentially as described (Bettinger et al., 2019) in order to calculate the ^18^O-met-containing to ^16^O-met-containing ratio for each methionine containing peptide feature in the MS1 spectra.

In cases where multiple methionine residues were mapped to a single peptide, the prevalence of alternative ^16^O/^18^O oxidized isotopic species were determined assuming a binomial distribution. For example, if a peptide contained 2 methionine residues, with *p* and *q* fractions of each methionine being in the ^16^O-oxidized form, then the relative fractions of ^16^O/^16^O, ^16^O/^18^O and ^18^O/^18^O isotopic variants of the peptides were expected to be present at relative fractions of *pq, p(1-q)+(1-p)q* and *(1-p)(1-q)*, respectively. In these scenarios, *p* and *q* were determined independently and were observed to always be within 0.1 fraction of each other. In other words, methionine residues that were contained in the same chymotryptic peptide, and hence were close to each another in sequence, always appeared to have similar oxidation rates. Nonetheless, in these scenarios, the different oxidation fractions were assigned to specific methionine residues within the peptide depending on the strength of evidence in the MS2 spectra indicating the presence of oxidation at a specific methionine (Cox and Mann, 2008). So in the example above, if *p* was greater *q*, and the MS2 spectrum of the ^16^O/^18^O peptide generated fragments that provided stronger support for the presence of ^16^O-oxidized form of the first methionine compared to the second methionine, *p* was assigned as the oxidized fraction of the first methionine, and *q* was assigned as the oxidized version of the second methionine. In cases where the strength of evidence was equally strong for both oxidized forms, was inconclusive, or MS2 spectra were not available for ^16^O-containing peptide forms, *p* and *q* were assigned to equal values that best fit the binomial distribution. For each methionine, the measured fractional oxidation levels from all corresponding peptide features were averaged to obtain a single consensus fractional oxidation measurement.

### Measurement of methionine protection factors in polymeric samples

We considered the following rate equation for oxidation of methionine by H_2_O_2_:

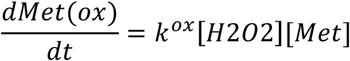

Where k^ox^ is the second order rate constant for methionine oxidation. Since H_2_O_2_ is present in excess, this is a pseudo-first order reaction where the experimentally observed first order rate constant (k^obs^) is:

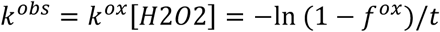

Where f^ox^ is the fraction of methionine residues that have been oxidized at a given H_2_O_2_ concentration and oxidation time (t). We measured k^ox^ of individual methionine residues by determining the slope of −ln (1 − *f*^*ox*^)/*t* as a function of [H_2_O_2_]. The rate constants were measured in monomers, hydrogels and liquid droplet samples. We observed that oxidation rate constants of methionine residues were slower in hydrogel and liquid droplets in comparison to monomeric samples at varying levels. The ratio of oxidation rate constants in the polymeric forms were calculated relative to the monomer to obtain “protection factors” for each methionine. Thus, protection factors provide a measure of how much methionine oxidation rates were impeded relative to monomers. For immuno-purified proteins oxidized in cells, oxidation rates for monomers are not available. Hence, the extent of oxidation is reported as a simple ratio of ^18^O-oxidized to ^16^O-oxidized peptides that has not been normalized relative to monomers.

## Acknowledgements

The authors thank Lillian Sutherland for extensive technical help and Deepak Nijhawan for encouragement and intellectual contributions. BPT was supported by NIH grant RO1-NS115546; SG by NIH grants R35-GM119502 and S10OD025242; SLM by NIH grant U01GM107632 as well as unrestricted funding provided by an anonymous donor.

